# Active vision during prey-capture in wild marmoset monkeys

**DOI:** 10.1101/2022.04.01.486794

**Authors:** Victoria Ngo, Julia C. Gorman, María Fernanda De la Fuente, Antonio Souto, Nicola Schiel, Cory T. Miller

**Author notes:** Equal contributions.

## Abstract

Here, we examined prey-capture in wild common marmosets (*Callithrix jacchus*) to explicate the active role of primate vision for high-precision, goal-directed motor actions in challenging natural environments. We observed distinct marmoset hunting strategies that each relied on the close integration of visual processes and dynamic biomechanical motor movements, but differed based on several factors including prey size/speed, substrate, and their relative distance to the prey. Marmoset positional behavior in these complex arboreal settings often involves extending the body, including inversions, while relying on vision to guide prey capture. Applying markerless pose estimation technology, analyses revealed details of how primate vision both guides and corrects rapid motor actions in real-time during prey-capture in the wild. These findings illustrate the significance of active vision during primate ethological behaviors in response to the very challenges the visual system evolved to solve.

## Summary

A foundational pressure in the evolution of all animals is the ability to travel through the world. Because movement relies on continuous streams of sensory information about the content and structure of the surrounding environment, sensory and motor systems are inherently coupled. While this relationship has been explored in several species^1–4^, it has been largely overlooked in nonhuman primates. The weight of our considerable knowledge about primate vision continues to rely on traditional paradigms in which restrained subjects view visual stimuli on screens^5^. Natural visual behaviors, by contrast, are typified by locomotion through the environment guided by active sensing as animals explore and interact with the world to gain different perspectives and integrate the resultant information^4,6^, a relationship well-illustrated by prey-capture^7–12^. During this natural behavior, individuals not only search for and track prey using vision, but must execute precise visually-guided motor plans for successful capture of prey while at the same time navigating the environment. Here, we sought to characterize prey-capture in wild marmoset monkeys (*Callithrix jacchus*) as they negotiated their dynamic, arboreal habitat to illustrate the inherent role of vision as an active process in natural nonhuman primate behavior. Marmosets are an ideal species in which to explore this issue because these monkeys both share the core properties of vision that typify the primate Order^13–18^, and are prolific hunters that prey on a diverse set of animals ranging from small insects to lizards^19–22^. We show the integral role of active vision in marmosets during the pursuit and capture of prey in several different contexts, including how marmosets positioned themselves in precarious arboreal settings to establish a clear visual field and effectively capture prey. Applying modern computer vision technologies for markerless tracking yielded novel findings pertaining to how marmosets track insects prior to initiating an attack and the rapid visually-guided corrections that guide hand movements during capture. These findings offer the first detailed insight into the active nature of vision to guide multiple facets of a natural goal-directed behavior in wild primates.

## Results

### [1] Prey-capture Strategies

Here we analyzed 288 4K UHD videos of marmosets engaged in active prey-pursuit and capture of insects in northeastern Brazil to characterize the relationship between natural, visual, and positional behavior in a wild primate. Hunting behaviors were grouped into three of the more typical tactics distinguished by the sequence of visually-guided motor actions employed in response to specific challenges of identifying and capturing different prey types in the dynamic forest environment^19^, though notably each prey-capture strategy was variable owing to the challenges of navigating the substrates. Although our analyses highlight hunting of insects, marmosets also actively hunt other moving prey (e.g., lizards, N = 27 videos).

Figure 1A illustrates ‘mouth capture’, the strategy employed by marmosets to track small, mobile prey with limited capture avoidance behaviors (e.g., ants, termites, etc., N = 80 Videos; Video S1) characterized by the overwhelming use of their mouth to capture insects than their hands (n = 73 observations; Mouth: 90.4%, Hands: 9.6%; p < 0.01, Fisher Exact Test). The prevalence of mouth use in this context is notable as it contrasts with all other prey-capture strategies observed in marmosets.

**Figure 1.**
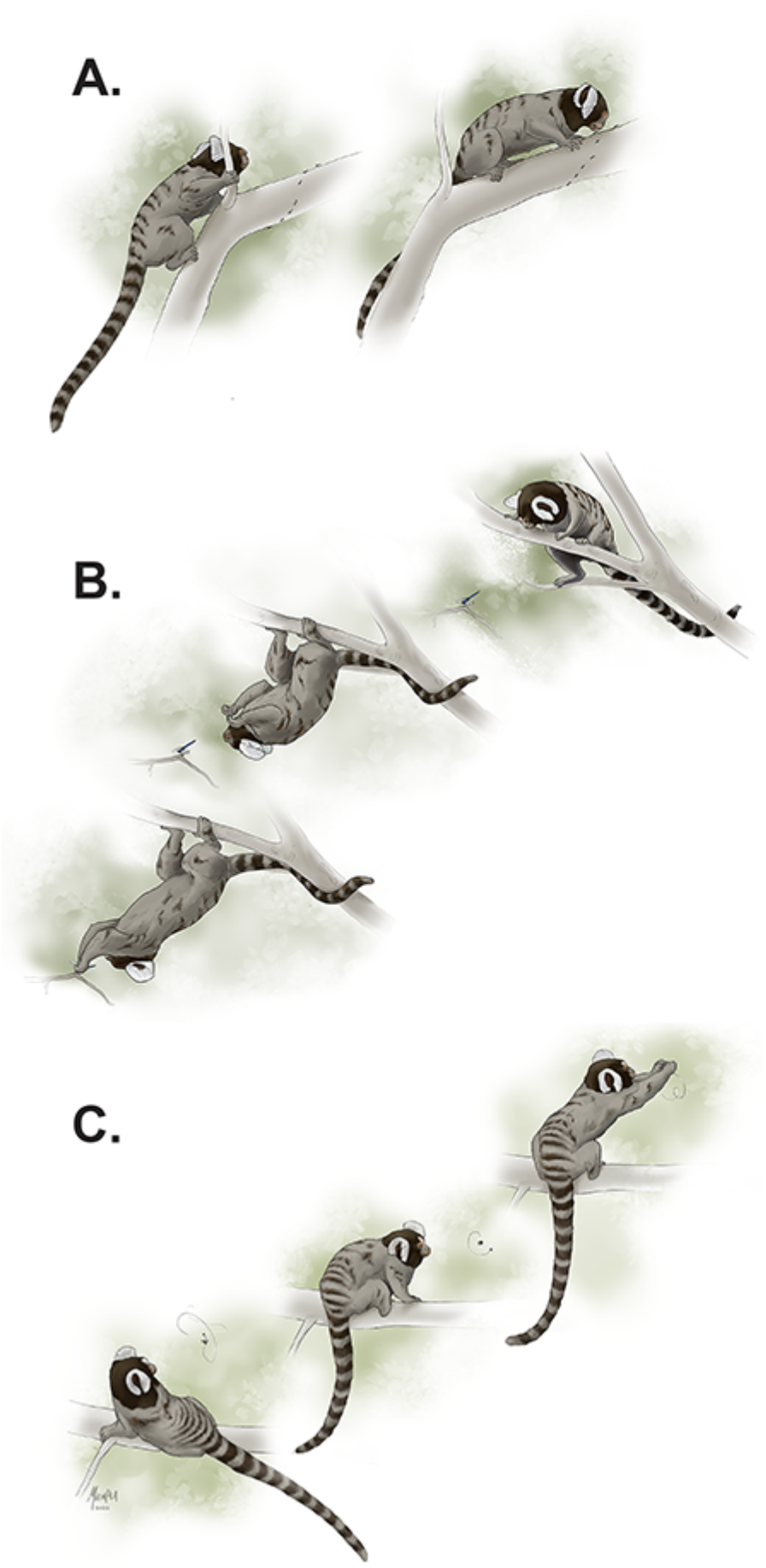
Marmoset Prey-Capture Strategies. (A) ‘Mouth Capture’; Marmosets typically capture small, non-evasive mobile prey (e.g., ants, termites, etc.) with their mouth. (B) ‘Stalk and Lunge’; Hunts of insects that can conceal their location through camouflage and also have effective predator avoidance strategies (e.g., stick bugs, grasshoppers, etc.) involve the marmoset slowly approaching or stalking the prey and optimizing the monkey’s positional behavior. Once positioned, marmosets typically pause for 1-5s immediately before initiating a ballistic hand grasp. (C) ‘Capture in Flight’; Prey-capture of flying insects requires precise visually guided motor planning and control, as well as adjustment of the arm trajectory in response to changes in the insect’s flight trajectory.

Figure 1B depicts ‘stalk and lunge’, the strategy employed by marmosets to capture stationary prey that often relied on camouflage to avoid capture (e.g., stick bugs, moths, grasshoppers, N = 93 videos; Video S2). Notably, insects that evolved mimicry increase the difficulty of their detection and are akin to a natural visual pop-out task to marmosets ^23,24^. Some of these species, while stationary when on the substrate, have fast evasive behaviors that monkeys must account for. Grasshoppers and dragonflies, for example, will rapidly jump or fly away if predators are detected, while stick bugs evade predation by holding their legs against their body and drop to the ground to hide among the leaves^25^. As a result, when hunting these prey, marmosets position themselves for the attack, sometimes slowly stalking the prey (Video S3). Once in position, marmosets typically pause for 1-5s before initiating a high-speed, ballistic grasp of the prey (Video S4).

Hunting flying insects is particularly challenging for marmosets with respect to both the demands on active vision and the role of sensory feedback to guide motor planning and control necessary for successful capture (Figure 1C; Video S5). In the recordings of marmosets pursuing flying beetles, their behavior typically abided by one of two strategies. It either involved a stationary monkey visually tracking the prey for a period of several seconds before initiating the ballistic grasp (N = 47 videos), or the animal simultaneously visually tracking and physically following the prey through the arboreal substrate as the insect’s flight pattern changed (N = 68 videos). Two-handed captures were more common for flying insects than one-handed or mouth captures (64.7%, N = 224 observations), likely because this tactic yielded a notably higher success rate (82.4%) than one-handed captures in wild marmosets (X^2^ (1) = 9.5, p = 0.002).

### [2] Positional Behavior during Arboreal Prey Capture

The stability and orientation of the substrate in which prey-pursuit occurred were significant factors that affected marmoset behavior before and during hunts. Marmosets countered these challenges with intricate changes in positional behavior that offered stability, while optimizing the likelihood of success. The use of adaptive positioning with multiple limbs on horizontal branches, including reaching both above (Figure 2A) and below (Figure 2B) the branch, and vertical branches that, likewise, involved pursuing prey above (Figure 2C) and below (Figure 2D) the substrate. Notably, one shared similarity across all positional behaviors during prey capture was optimizing visual access of the target while balancing the need for stability, so as to avoid falling. This dynamic is well illustrated by the manner in which marmosets extended their body at various angles from the more stable arboreal substrates in pursuit of insects that were often found on small unstable branches. In these settings, the monkeys often extended their body using only their hind limbs as support at a range of angles from the substrate for successful prey capture. We quantified these positional behaviors by measuring the monkey’s angle of attack and the extent to which individuals extended their body in the attack (Figure S1). The average percent body length extension during prey capture for marmosets on top of the substrate was 63.8% ± 54.5% (mean ± SD), while body extension when clinging under the substrate was 65.7% ± 36.4% (mean ± SD). Marmosets hunting on a vertical substrate with the head pointing upwards extended their body 51.2% ± 38.8% (mean ± SD), while marmosets hunting on a vertical substrate with the head pointing downwards extended their body 57.3% ± 29.1% (mean ± SD) (Figure 2E). When marmosets are under a horizontal substrate, they will typically extend downwards 95% of the time (n = 20/21, N = 158 observations) as they are already hanging from the branch with their lower limbs, whereas when they are on top of the horizontal substrate, they will extend downwards only about 12% of the time (n = 9/73, N = 158 observations). These results illustrate the flexible nature of marmoset prey capture in arboreal environments that reflect the need to balance and optimize stable positioning and vision for successful prey capture.

**Figure 2.**
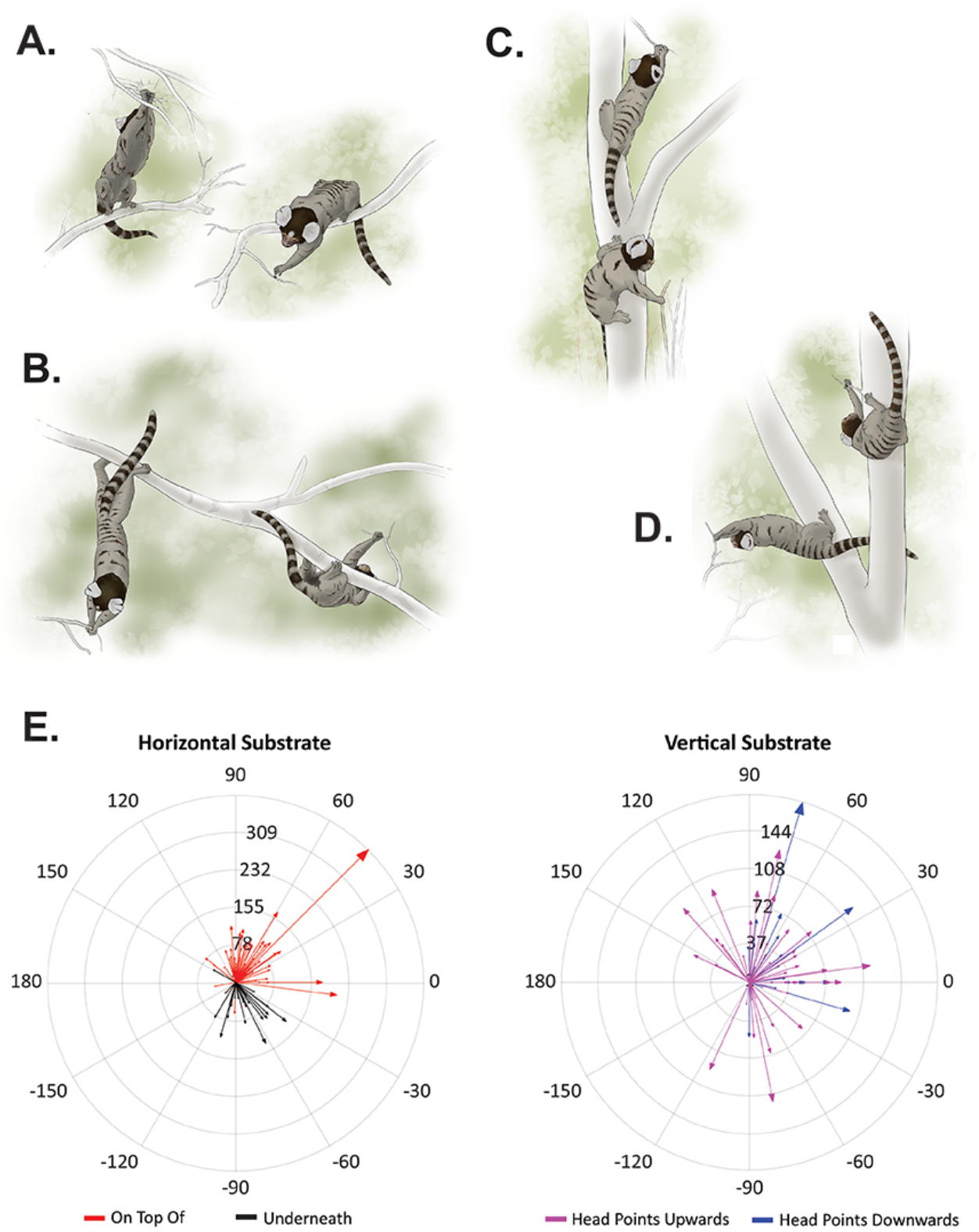
Positional Behavior Preferences Based on Substrate Orientation. (A-D) Illustrations depicting the range of positional behaviors during hunting (reaching above or below) while the marmoset is situated (A) on horizontal substrate, (B) under horizontal substrate, (C) on vertical substrate with the normal vector of the marmoset’s head pointing up, and (D) on vertical substrate with the normal vector of the marmoset’s head is pointing down. (E) The polar plots represent the marmoset’s angle of extension when reaching for prey depending on the substrate being grasped and percent change in body length of extension calculated in pixel units. Arrows pointing below the horizontal axis refer to marmosets reaching downwards while hunting. Horizontal substrates are shown to the left and vertical substrates to the right.

### [3] Gaze-Tracking of Flying Insects

To precisely quantify visual behaviors during prey-capture in wild marmosets, we next applied the markerless tracking technology SLEAP^26,27^ to a subset of videos that met a set of criteria related to video recording quality and modeling accuracy (See Methods). These analyses focused on hunts of flying insects because this context highlights the unique challenge of successfully capturing a moving insect in three-dimensional space and the role of vision as an active process for a goal-directed, high-precision motor action. We distinguished between two phases of the marmoset hunt for these analyses based on the visual challenges. In the first phase, marmosets visually track the prey and, once identified, plan for the attack. During the second phase, marmosets initiate the ballistic grasp based on the anticipated location of the prey.

We first analyzed head movements in the final seconds prior to the initiation of the ballistic grasp as a proxy for gaze-tracking, including how they covaried with the flight path of the insect prey. Frame grabs from an exemplar video show three time points over 500ms immediately before the ballistic grasp is initiated and highlight the close relationship between head and insect movements that occur during this time period (Figure 3A). However, this coupling was largely limited to the final period, 1.5s, just before initiating capture. Figure 3B shows the change in XY coordinates for the head and insect movements in the same exemplar video over a longer period of time. The changes in flight trajectory occur early in the video, but only exhibit a similarity over the final second. This general observation was also evident when analyzing all videos that reached our criteria. Namely, the correlation between marmoset head and insect movements was highly variable until the final 1.5s prior to initiating the ballistic hand movement for capture. The Pearson correlation coefficient between head and insect movements remained between 0.5 - 1 (N = 5) for the entirety of this time period (Figure 3C). One other notable result from this analysis was the prodigious increase in head velocity over the 500ms prior to initiating the capture action (Figure 3D). This likely suggests that marmosets were closely tracking the flight path, potentially reflecting a change in the attentional demands necessary for accurate motor planning and successful execution of the capture. This analysis may also suggest that marmosets have a preference for a particular type of insect flight behavior when initiating an attack, when the insects are moving along a continuous path rather than at times when the path was less predictable.

**Figure 3.**
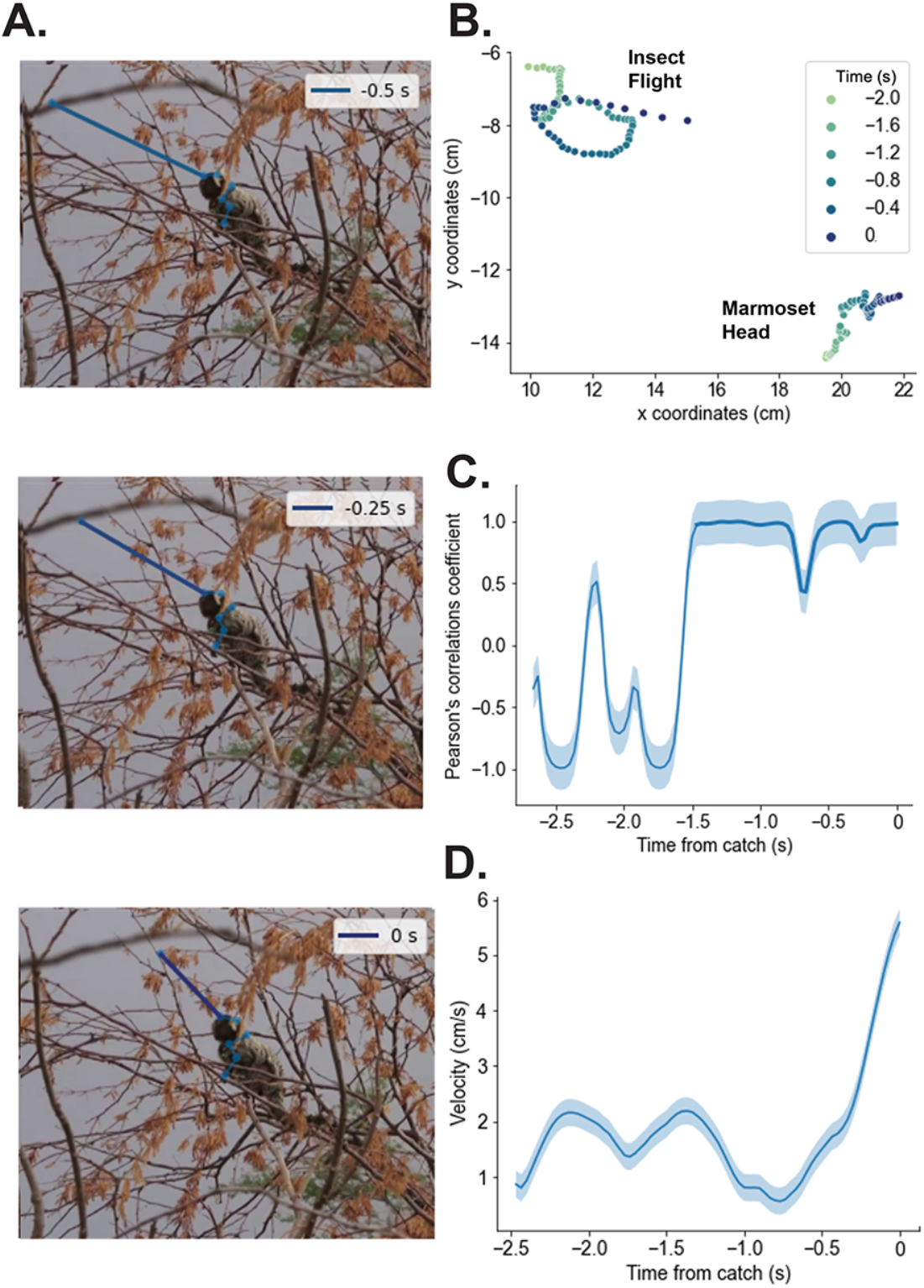
Gaze tracking insects during prey-pursuit. Marmoset gaze-tracking of flying insects is characterized by the relationship between marmoset head and insect movements prior to initiating capture. (A) Three single frames from the final 500ms before prey capture illustrate coordinated movements between the marmoset head and insect flight trajectory in an exemplar video. (B) Raw XY positional data of both the marmoset’s head movement (bottom right) and insect’s flight pattern (top left) from the example screen shown in A over the 2s prior to initiating the ballistic grasp for prey capture. (C) Pearson’s correlation coefficient between the marmoset head and insect flight in a sliding window over the 2.5s before prey capture was initiated. Zero (0) indicates the onset of the ballistic grasp. 95% Confidence Intervals are shown in shading. (D) Average velocity (cm/s) of the marmoset head over the same time period as in C. 95% Confidence Intervals are shown in shading.

### [4] Visually-Guided Prey Capture

Because the position of flying insects is constantly changing, successful capture relies on a complement of two overlapping visually-guided processes. First, marmosets must anticipate the likely position of the insect in three-dimensional space at a point in time. Second, as insects frequently change their flight pattern, marmosets need to make quick adjustments in response to changes in insect trajectory after the ballistic grasp is initiated. To quantify the second phase of the hunt, visually-guided prey-capture, we again applied SLEAP to a subset of videos that met our criteria for these analyses (See Methods). Figure 4A depicts a parallel series of frame grabs from an exemplar video and the respective XY coordinates of the hand movements when a marmoset reaches for and grasps a flying insect that illustrate how the respective position of the hands and insect change over this time. The overall changes of these hand movements over the duration of the event are shown in contrast to the optimal trajectory in Figure 4B for this exemplar video. Importantly, the hands did not follow this optimal trajectory in any of the videos analyzed, but rather diverged in several quantifiable ways. First, the hand movements had several inflections (i.e., or points at which sudden opposing adjustments were made) reflecting the changes in trajectory during the motor action (Avg = 2.4, Range = 0-7; Figure 4C). Second, the average tortuosity - i.e. the length of the movement including the turns/bends - for the hand movement trajectory was 3.08, which significantly diverged from the optimal path (t(24) = 5.31, p < 0.0001; Figure 4D). These corrections in hand trajectory are occurring at high speeds, as the latency to peak velocity occurred at an average of 224.44ms (Figure 4E), while the mean duration of the entire ballistic grasp during capture was 375.61ms (Figure 4F). These analyses indicate that marmosets made real-time visually-guided corrections to the path of the hand trajectory during the short interval of time from the initiation of the motor action until the insect was captured, likely as a result of changes in the flight path of the prey during that interval. This finding shows that, like other human and nonhuman primates^28–30^, wild marmoset hand movements when targeting the capture of flying beetles are under continuous visual control, and can be modified through feedback at different points over the motor actions, in response to the exact types of natural challenges the visual system evolved mechanisms to overcome.

**Figure 4.**
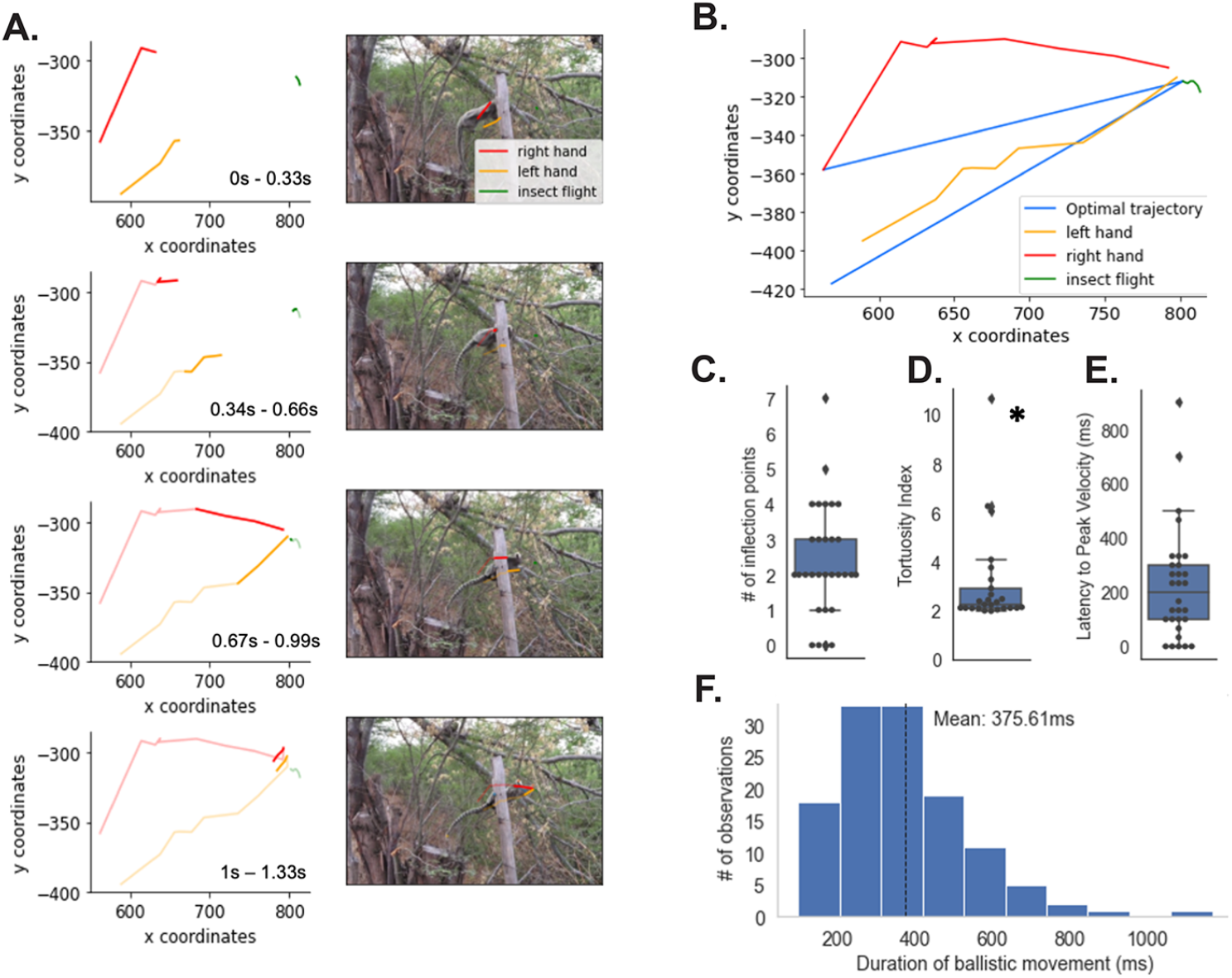
Visually-guided prey-capture. (A) Left plots the XY coordinates of the right and left hands of the marmoset and insect at four different time points during an exemplar video. The corresponding individual frames are shown to the right. (B) Summary of A shows the total path of both hands and the insect, as well as the optimal trajectory in blue for each hand. (C) Number of inflection points during reaches from all videos analyzed. (D) Plots the tortuosity index for all prey-capture reaches. (E) Latency to peak velocity (ms) during ballistic grasps for flying prey. (F) Duration of ballistic movement (ms) from initiation of the motor action to successful grasp of the prey.

## Conclusions

Here we demonstrate that active vision is integral to successful prey-capture in wild marmosets. The dynamic biomechanical movements these monkeys take through the arboreal landscape when searching for living prey, reflect active choices to acquire the diverse visual perspectives that optimize the different behavioral strategies used in each instance for an effective capture (Figure 1). Owing to the dynamic nature of the arboreal landscape and the often precarious locations that insects were found, marmosets were routinely confronted with unique challenges on how to position themselves in order to successfully grasp the insects once they were recognized. Far from being a passive process, vision is closely coupled to all elements of prey capture in marmosets, similarly to other species^7–11^. A compelling advantage of this natural behavior is that it comprises of visual processes studied independently in primate vision for decades - discrimination, recognition, motor planning, decision-making, and visually-guided selection, amongst others - within a single, cohesive visual behavior^34^. These visual processes are in fact not separable, but operations within an integrated behavioral sequence that is representative of the distinct challenges that together have driven the evolution of the complementary mechanisms that typify the primate visual system. Far from being stereotyped, marmoset prey-capture was remarkably flexible, likely reflecting the need to be adaptable to the immediate environment for successful predation. This feature further underscores the advantages of prey-capture for understanding the factors affecting problem solving and cognitive control in primates under variable natural conditions. As not all theoretical models of vision developed in more traditional laboratory settings are likely reflective of how this system functions under real-world scenarios^31^, natural primate behaviors such as prey-capture will likely be necessary to elucidate remaining key questions about primate brain function.

## Acknowledgements

We thank Drs. Talmo Perreira, Adam Fishbein, Dori Grijseels, and Arthur Lefevre for comments on the manuscript, Dr. Geraldo Baracuhy for allowing us to conduct this research at Baracuhy Biological Field Station and Sarah Mientka for the marmoset illustrations. This work was supported by grants from the AFOSR (19RT0316) and NIH (1UF1NS116377) to CTM.

## Methods

### Study Site and Subjects

High-resolution videos were recorded of wild marmosets inhabiting the semiarid scrub-forests in the Baracuhy Biological Field Station in Northeast Brazil (7°31’42”S, 36°17’50”W)^32,33^. Data collection was conducted between March 2020 and June 2021 following two social groups-House Group (mean group size of 9 animals) and Coqueiro Group (mean group size of 8 animals).

### Data Analysis

Analyses were performed on the 288 Ultra HD video recordings (4K; width: 3840 pixels; height: 2160 pixels) of marmoset prey-capture on the Sony FDR AX-53 camcorder at a frame rate of 29.970 fps. Videos may contain one or more observations of a hunting strategy by one or more subjects being filmed. 287 videos that were not reflected in the final analyses exhibit marmosets foraging for food by leaf manipulation or opportunistic prey capture^19^. Videos recorded in the field were uploaded in Brazil onto a shared server and cataloged at UCSD.

#### Positional behavior during arboreal prey capture

Quantitative analysis shown in Figure 2 was performed using Adobe Premiere Pro version 22.0 and Adobe Photoshop version 22.5.1. Premiere was used to quantify the duration of a subject’s hunting action frame by frame up to hundredths of a second, including the motor planning pause before a ballistic movement, and the start to completion of a ballistic movement based on the type of prey the marmoset went after. Using Premiere, multiple instances of gaze tracking were recorded, whether it was directed towards the prey or when it shifted away. The start of the gaze was defined as when the marmoset’s head movement stops scanning and the head either moves in closer or cocks from side to side before the body starts to move in the same direction. The end of the gaze period is defined when the marmoset’s head movement shifts away from the targeted prey as it moves closer, usually to redirect its attention to the route they are trying to take or to check if there are other competitors around. The gaze can then resume when the head movement turns back to the direction of where the target is/where the target is moving. Frames depicting a side profile of the moment before and after a ballistic movement was achieved were exported into Adobe Photoshop for further examination.

Since body lengths of animals in the wild were not possible, the ruler tool in Adobe Photoshop was used to obtain measurements in pixels including the marmoset’s body length pre-and post-ballistic movement in exported frames, allowing us to attain the percent length of extension. For the pre-ballistic measurement of the marmoset’s body, it was measured from the base of the tail or foot to the top of their forehead or mouth, depending on the visibility of each body part in the frame. The post-ballistic measurement was measured from either the same tail base or foot—if visible and in instances where they are using their legs to extend—to the extended body part being used to grab (either the extended hand(s) or mouth). Additionally, the ruler tool was used to estimate the angle of extension from the substrate a subject opts for depending on how they are oriented on that substrate. The angle was measured from the parallel substrate, to the base of where the marmoset is clinging onto, and to the extended arms. Owing to the limitations of a 2D video, videos demonstrating body extension along the Z plane were not taken into consideration for measuring to minimize the chance of error.

The polar plot in Figure 2E was created using a custom MATLAB script to demonstrate the marmoset’s angle of extension depending on the substrate they are hunting on (horizontal or vertical) and the length of the arrow depicts a visual representation of the percent body length of extension in pixels. The “-” sign refers to the downward direction the marmoset is reaching in (Figure 2E). To obtain the normalized percent change in length of extension, the following formula was used:

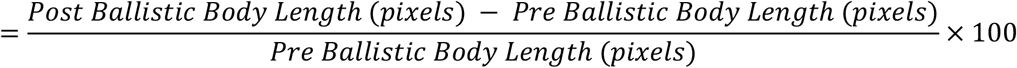

Since only one camera angle was often available for analysis, the ability to measure velocity during a capture was not feasible. Instead, using the aforementioned formula, we examined the normalized percent change in the marmoset’s body length over the duration of the ballistic movement during a hunting sequence to capture their change in motion.

#### Markerless tracking analyses

To more precisely quantify the marmoset’s visuomotor integration that occurs when hunting, we employed computer vision technology for markerless tracking on one dynamic hunting strategy involving a one- or two-handed capture mid-air following rapid head-gaze tracking behavior of flying beetles (Coleoptera). Down-sampled videos (1280 × 720) were uploaded onto SLEAP (Social LEAP Estimates Animal Poses) ^26,27^ where deep learning was employed for markerless tracking of the marmoset body, whereas the flying insect was hand annotated either on a separate project or following the inference process. The right hand, left hand, middle top of head, left ear tuft, and right ear tuft of the marmoset in the video were manually annotated in a small percentage (∼15%) of frames. Then, active learning takes place as we train a neural network through the single animal pipeline to estimate positions of the marmosets body parts by running inference. While pose estimation was computed for the entire video clip (typically ∼15s) to improve the robustness and accuracy of the model, analyses presented here focused on the few seconds before and during insect capture. These select frames were further refined for analysis by additional hand annotations. H5 files were then exported for use in additional analyses.

##### Gaze-tracking flying beetles

Analysis was performed on videos that met the following criteria: (1) Video maintained stable, continuous focus of both the insect and the animals head for at least 2 seconds prior to initiating capture, (2) the animal was not chasing the insect during this period or otherwise locomoting, but remained sitting on the substrate and (3) there were no obstructions of either the insect or marmoset that would affect annotation of the video. Five videos met these strict criteria and were used in the analysis typically due to the flying insect not remaining in focus for sufficient periods of time during the video. The frame number of both the start of the marmoset gaze and the end of the gaze/start of the ballistic movement was marked in each video. These frame numbers were validated by two other lab members. Videos were processed in Python 3.6 and the aforementioned time periods mentioned during the gaze period were analyzed (Figure 3A). Gaze estimates in 3B were made between the SLEAP label that demarcated the middle of the head and the SLEAP bug label. A line was drawn from the two to show an approximate gaze line. In 3C, velocity of both the head SLEAP labels and the insect’s SLEAP label were calculated. A rolling correlation was taken to quantify the relationship between the head and the bug in the final second. The Python pandas package outputs the Pearson’s Correlation Coefficient in a sliding bin window with a bin size of 10. In order to characterize the period of high correlation before the bug catch, a rolling average of bin size 10 was taken and any correlation coefficient above 0.5 was considered significant. A 95% confidence interval was calculated using Python in order to show the variability of the data in both 3C and 3D.

##### Visually-guided prey capture

These analyses were performed on videos that maintained stable, continuous focus and/or accurate model prediction of both the insect and at least one of the animal’s hands for the duration of time from the initiation of the ballistic grasp till prey capture. Analyses were performed on 15 videos that met these criteria and had the least occlusion by vegetation. Lines were drawn from the start of the ballistic movement to the point of prey capture to demonstrate the most direct path - the optimal trajectory - of a reach if the insect’s flight path was perfectly predicted. We next applied two different measurements to quantify how much marmoset hand trajectories deviated from the optimal trajectory: ‘Number of Inflection points’ and ‘Tortuosity’. The left and right hands were analyzed separately in each video because both hands were not always clearly visible. This yielded 15 left hand events and 9 right hand events from the videos used in analyses here.

- *Inflection points*. This analysis identified the number of instances that marmosets modified the direction of their hand movement during the ballistic grasp leading to prey capture. This measure was taken by first calculating the distances from the ideal line and then calculating the number of local maxima and minima. Values were validated by marking the point at which a local max and min were found.
- *Tortuosity*. To better quantify the changes and curvature of the arm reaches during prey capture of flying insects, we measured the tortuosity of the reach trajectory. Tortuosity was calculated to show the ratio between the distance of the optimal trajectory or total length (L) and the actual movement of the hand or the path length (C). This ration was then 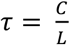. In order to test for significance, a paired t-test was performed against the null hypothesis of 1, since 1 would be the value of τif *C* = *L*.
- *Latency to peak velocity*. We calculated the velocity of the hand over the course of the ballistic action to determine the time at which the velocity was at its maximum.

To calculate the duration of ballistic reaches to capture flying insects for Figure 4E, we lowered our selection criteria to all videos in which the body was visible (n = 123 observations). The frame numbers were denoted for the duration of the ballistic movement and the average was calculated.

## Supplemental Materials

**Figure S1.**
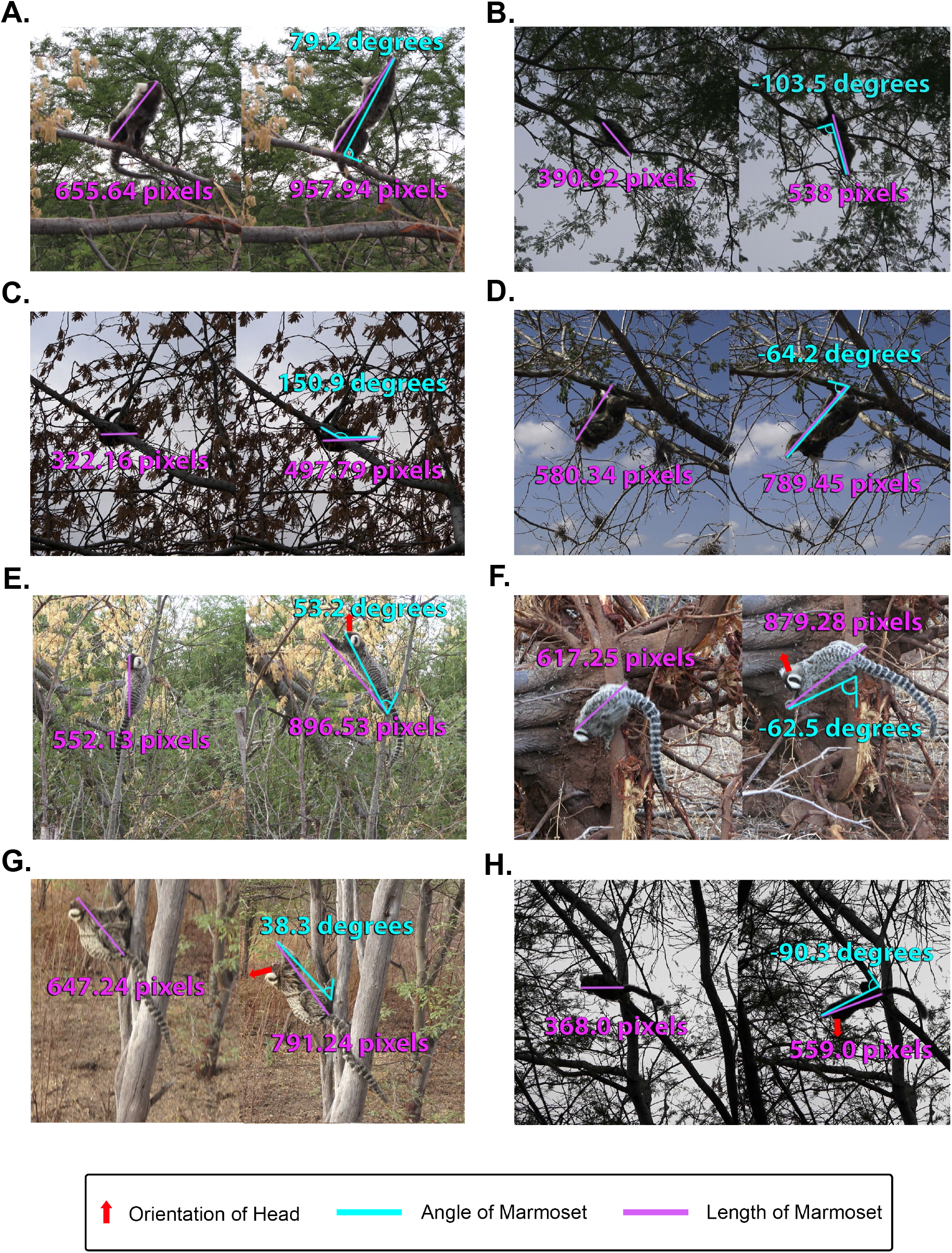
Measurements taken from Adobe Photoshop of body length pre- and post-ballistic movement, to calculate percent change in length of extension, and the angle the marmoset extends relative to the substrate. Positive angles refer to when marmosets are reaching upwards (A, C, E, G) and negative angles refer to when they reach in a downward direction (B, D, F, H). Quantitative data taken from examples of the extensive range of marmoset positional behaviors including: on top of horizontal substrate (A, B), under horizontal substrate (C, D), on vertical substrate with normal vector of marmoset’s head pointing up (E, F), and on vertical substrate with normal vector of marmoset’s head pointing down (G, H). Related to Figure 2.

<u>Video S1</u>

Adult female marmoset follows a scattered trail of ants along the tree trunk/branches and uses its mouth as the primary approach of hunting. Related to Figure 1A.

<u>Video S2</u>

Juvenile marmoset spots camouflaged cricket in a stationary position and repositions before performing a high-speed ballistic grasp with both hands. Related to Figure 1B.

<u>Video S3</u>

Adult female marmoset slowly stalks the dragonfly by dragging its body along the tree branch and pauses for 1.25s before achieving an upper body lunge with both hands. Related to Figure 1B.

<u>Video S4</u>

Adult male marmoset pauses for several seconds before executing a full body lunge directly upwards to capture stick bug blending among the thin branches using both hands. Related to Figure 1B.

<u>Video S5</u>

Adult male marmoset surrounded by several flying beetles while in a stationary position. Visually tracks multiple beetles as they fly into marmoset’s field of view and attempts several times to capture them using one or two hands. Related to Figure 1C.

All supplemental videos are available here: https://drive.google.com/drive/folders/1o1ww0krcILuK_oHqABEtIk5g7ALp8ZAy?usp=sharing

## References

1. Kleinfeld, D., Ahissar, E., and Diamond, M.E. (2006). Active sensation: insights from the rodent vibrissa sensorimotor system. Curr. Opin. Neurobiol. 16, 435–444.

2. Parker, P.R.L., Brown, M.A., Smear, M.C., and Niell, C.M. (2020). Movement-Related Signals in Sensory Areas: Roles in Natural Behavior. Trends Neurosci. 43, 581–595.

3. Wachowiak, M. (2011). All in a Sniff: Olfaction as a Model for Active Sensing. Neuron 71, 962–973.

4. Gibson, E.J. (1979). The ecological approach to visual perception (Houghton Mifflin).

5. Leopold, D.A., and Park, S.H. (2020). Studying the visual brain in its natural rhythm. Neuroimage 216, 116790.

6. Schroeder, C.E., Wilson, D.A., Radman, T., Scharfman, H., and Lakatos, P. (2010). Dynamics of Active Sensing and perceptual selection. Curr. Opin. Neurobiol. 20, 172–176.

7. Palleroni, A., Miller, C.T., Hauser, M., and Marler, P. (2005). Predation: Prey plumage adaptation against falcon attack. Nature 434, 973–974.

8. Lin, H.-T., and Leonardo, A. (2017). Heuristic Rules Underlying Dragonfly Prey Selection and Interception. Curr. Biol. 27, 1124–1137.

9. Mischiati, M., Lin, H.-T., Herold, P., Imler, E., Olberg, R., and Leonardo, A. (2015). Internal models direct dragonfly interception steering. Nature 517, 333–338.

10. Michaiel, A.M., Abe, E.T., and Niell, C.M. (2020). Dynamics of gaze control during prey capture in freely moving mice. Elife 9.

11. Wagner, H., Kettler, L., Orlowski, J., and Tellers, P. (2013). Neuroethology of prey capture in the barn owl (Tyto alba L.). J. Physiol. Paris 107, 51–61.

12. Hoy, J.L., Yavorska, I., Wehr, M., and Niell, C.M. (2016). Vision Drives Accurate Approach Behavior during Prey Capture in Laboratory Mice. Curr. Biol. 26, 3046–3052.

13. Mitchell, J.F., Reynolds, J.H., and Miller, C.T. (2014). Active vision in marmosets: a model system for visual neuroscience. J. Neurosci. 34, 1183–1194.

14. Mitchell, J.F., and Leopold, D.A. (2015). The marmoset monkey as a model for visual neuroscience. Neurosci. Res. 93, 20–46.

15. Mitchell, J.F., Priebe, N.J., and Miller, C.T. (2015). Motion dependence of smooth pursuit eye movements in the marmoset. J. Neurophysiol. 113, 3954–3960.

16. Coop, S.H., Yates, J.L., and Mitchell, J.F. (2021). Foveal remapping of motion in area MT of the marmoset monkey. J. Vis. 21, 2638–2638.

17. Hori, Y., Cléry, J.C., Selvanayagam, J., Schaeffer, D.J., Johnston, K.D., Menon, R.S., and Everling, S. (2021). Interspecies activation correlations reveal functional correspondences between marmoset and human brain areas. Proc. Natl. Acad. Sci. U. S. A. 118.

18. Hung, C.C., Yen, C.C., Ciuchta, J.L., Papoti, D., Bock, N.A., Leopold, D.A., and Silva, A.C. (2015). Functional mapping of face-selective regions in the extrastriate visual cortex of the marmoset. J. Neurosci. 35, 1160–1172.

19. Schiel, N., Souto, A., Huber, L., and Bezerra, B.M. (2010). Hunting strategies in wild common marmosets are prey and age dependent. Am. J. Primatol. 71, 1–8.

20. Schiel, N., and Souto, A. (2017). The common marmoset: An overview of its natural history, ecology and behavior. Dev. Neurobiol. 77, 244–262.

21. Digby, L., and Barreto, C.E. (1998). Vertebrate predation in common marmosets. Neotrop. Primates 6, 124–126.

22. Souto, A., Bezerra, B.M., Schiel, N., and Huber, L. (2007). Saltatory search in free-living Callithrix jacchus: Environmental and Age influences. Int. J. Primatol. 28, 881–893.

23. Bichot, N.P., and Schall, J.D. (1999). Saccade target selection in macaque during feature and conjunction visual search. Vis. Neurosci. 16, 81–89.

24. Wang, Q., Cavanagh, P., and Green, M. (1994). Familiarity and pop-out in visual search. Percept. Psychophys. 56, 495–500.

25. Markle, S. (2008). Stick Insects: Masters of Defense (Lerner Publications).

26. Pereira, T.D., Aldarondo, D.E., Willmore, L., Kislin, M., Wang, S.S.H., Murthy, M., and Shaevitz, J.W. (2019). Fast animal pose estimation using deep neural networks. Nat. Methods 16, 117–125.

27. Pereira, T.D., Shaevitz, J.W., and Murthy, M. (2020). Quantifying behavior to understand the brain. Nat. Neurosci. 23, 1537–1549.

28. Oostwoud Wijdenes, L., Brenner, E., and Smeets, J.B.J. (2011). Fast and fine-tuned corrections when the target of a hand movement is displaced. Exp. Brain Res. 214, 453– 462.

29. Goodale, M.A., Pelisson, D., and Prablanc, C. (1986). Large adjustments in visually guided reaching do not depend on vision of the hand or perception of target displacement. Nature 320, 748–750.

30. Song, J.-H., Takahashi, N., and McPeek, R.M. (2008). Target selection for visually guided reaching in macaque. J. Neurophysiol. 99, 14–24.

31. Matthis, J.S., Muller, K.S., Bonnen, K.L., and Hayhoe, M.M. (2022). Retinal optic flow during natural locomotion. PLoS Comput. Biol. 18, e1009575.

32. De la Fuente, M.F., Sueur, C., Garber, P.A., Bicca-Marques, J.C., Souto, A., and Schiel, N. (2022). Foraging networks and social tolerance in a cooperatively breeding primate (Callithrix jacchus). J. Anim. Ecol. 91, 138–153.

33. Caselli, C., Ayres, P.H.B., Castro, S., Souto, A., Schiel, N., and Miller, C.T. (2018). The role of extra-group encounters in a Neotropical cooperative breeding primate, common marmosets: Field playback experiments. Anim. Behav. 136, 137–146.

34. Miller CT.; Gire D; Hoke K; Huk A; Kelley D; Leopold D; Smear M; Theunnisen F; Yartsev M; and Niell C (2022) The Language of the Brain is Natural Behavior. Current Biology, In Press

